# A novel soluble ACE2 protein totally protects from lethal disease caused by SARS-CoV-2 infection

**DOI:** 10.1101/2021.03.12.435191

**Authors:** Luise Hassler, Jan Wysocki, Ian Gelarden, Anastasia Tomatsidou, Haley Gula, Vlad Nicoleascu, Glenn Randall, Jack Henkin, Anjana Yeldandi, Daniel Batlle

**Author notes:** **Correspondence:** Dr. Daniel Batlle, Division of Nephrology and Hypertension, Department of Medicine, Northwestern University, The Feinberg School of Medicine, 320 E Superior, Tarry 14-727, Chicago, IL 60611.

## Abstract

Severe acute respiratory syndrome coronavirus type 2 (SARS-CoV-2) uses full-length angiotensin converting enzyme 2 (ACE2), which is membrane bound, as its initial cell contact receptor preceding viral entry. Here we report a human soluble ACE2 variant fused with a 5kD albumin binding domain (ABD) and bridged via a dimerization motif hinge-like 4-cysteine dodecapeptide, which we term ACE2 1-618-DDC-ABD. This protein is enzymatically active, has increased duration of action in vivo conferred by the ABD-tag, and displays 20-30-fold higher binding affinity to the SARS-CoV-2 receptor binding domain than its des-DDC monomeric form (ACE2 1-618-ABD) due to DDC-linked dimerization. ACE2 1-618-DDC-ABD was administered for 3 consecutive days to transgenic k18-hACE2 mice, a model that develops lethal SARS-CoV-2 infection, to evaluate the preclinical preventative/ therapeutic value for COVID-19. Mice treated with ACE2 1-618-DDC-ABD developed a mild to moderate disease for the first few days assessed by a clinical score and modest weight loss. The untreated control animals, by contrast, became severely ill and had to be sacrificed by day 6/7 and lung histology revealed extensive pulmonary alveolar hemorrhage and mononuclear infiltrates. At 6 days, mortality was totally prevented in the treated group, lung histopathology was improved and viral titers markedly reduced. This demonstrates for the first time in vivo the preventative/ therapeutic potential of a novel soluble ACE2 protein in a preclinical animal model.

## Introduction

Severe acute respiratory syndrome coronavirus type 2 (SARS-CoV-2), like SARS-CoV and other selected coronavirus, uses angiotensin converting enzyme 2 (ACE2) as its main receptor (1–8). After binding of the SARS-CoV-2 S spike to cell membrane-bound ACE2 there is priming by proteases, namely TMPRSS2, that are needed for the fusion and internalization of the ACE2-viral spike complex (1, 9–11). As early as 2003, when ACE2 was discovered to be the host receptor for SARS-CoV, in vitro experiments with native soluble ACE2 showed that this protein can neutralize the S spike protein of SARS-CoV (2, 3). In keeping with this concept early in 2020, we hypothesized that soluble ACE2, by acting as a decoy, could be used as a potential therapeutic approach for SARS-CoV-2 infection (12). There have been recent *in vitro* studies testing decoys for SARS-CoV-2 (13–20). *In vivo* studies, however, in suitable animal models, to our knowledge, are lacking. Monteil *et al* first showed that native soluble ACE2 can neutralize SARS-CoV-2 infectivity in human organoids (19). This soluble protein, moreover, is currently on a clinical trial (NCT00886353) where it is given twice a day intravenously to seriously ill COVID-19 patients. There is also a report of a patient that was given this protein for compassionate use (21). This patient survived, but one cannot be sure that the therapy was directly responsible for the good outcome.

Our lab has bioengineered soluble ACE2 variants that are shorter than native soluble ACE2, and therefore are more suitable for treatment of kidney disease as they are amenable for glomerular filtration (22). We have recently generated a short human ACE2 variant that has 618 amino acids, ACE2 1-618 and exhibits high enzymatic activity (16). This short ACE2 variant, moreover, was fused with a small (5-kD) albumin binding domain (ABD) as a way to extend *in vivo* duration of action (16). The duration of action of this novel protein assessed by plasma enzymatic activity is about 3 days whereas the duration of action of native soluble ACE2 and naked ACE2 1-618 is about 8 hours (16, 23). We have recently reported that all these proteins, ACE2 1-618, ACE2 1-740 and ACE2 1–618-ABD, can neutralize SARS-CoV-2 infectivity in human kidney organoids (16).

Key properties of an optimal soluble ACE2 protein for COVID-19 use should include strong binding for the receptor binding domain (RBD) of SARS-CoV-2 and enhanced duration of action. ACE2 1-618 and ACE2 1-618-ABD are monomers. Viral spike trimers may be more amenable to simultaneous binding to dimeric ACE2 forms analogously to that found with enhanced binding of dimeric diabodies to trimeric fusion protein in RSV (24). We therefore reasoned that a dimerized ACE2 1-618-ABD with extended duration of action might display increased affinity to the S1 spike of SARS-CoV-2. To accomplish dimerization, ACE2 1-618-ABD was bridged by fusion with a hinge-like 13-mer KCHWECRGCRLVC (DDC). Here we report the generation of the chimera termed ACE2 1-618-DDC-ABD and demonstrate its characteristics: 1) enhanced duration of action; 2) increased binding to SARS-CoV-2; and 3) full enzymatic activity. Since rodents are resistant to SARS-CoV-2 infection, we used a transgenic mouse model rendered susceptible to infection by introduction of the human ACE2 protein, the k18-hACE2 mice (25). In this model, inoculation with SARS-CoV and SARS-CoV-2 causes a lethal disease (25–30). Here we report that administration of a novel soluble ACE2 protein to these mice prevented the lethality observed in untreated animals altogether and resulted in a mild disease with reversible lung damage.

## Methods

### Generation of Human ACE2 1-618-DDC-ABD protein

Recombinant human ACE2 protein chimeras were generated using an approach similar to that recently described (16, 22). Briefly, a cDNA coding for C-terminal portion of the 618 amino acids fragment of human ACE2 protein (termed *ace2 1–618*) was fused with the *abd* cDNA encoding for a small ABD protein (5-kD) using a flexible linker (*g4s3*) placed between the N terminal end of the *abd* cDNA (IDT) and the C-terminal end of *ace2 1-618*. This resulted in prolonged *in vivo* duration of action of ACE2 as described previously (16). To achieve dimerization of the ACE2-ABD chimeric protein we inserted cDNA coding for a hinge-like region containing a dodecapeptide motif (31)termed DDC between *g4s3* and the c-terminus of the cDNA for *ace2 1-618*. Thus, 20 additional amino acids intervene between the C terminus of ACE2 1-618 and the N terminus of ABD. The cDNA of the fusion chimera termed *ace2 1–618-ddc-abd* was then inserted into pcDNA3-4 plasmid (Invitrogen) using custom synthesized complementary primers (IDT) and the Gibson assembly kit (NEB). After verifying the DNA sequence of the pcDNA3-4 fused with the cDNA for *ace2 1-618-ddc-abd*, the protein was produced in expi293 cells and purified using Fast Protein Liquid Chromatography on Q-Sepharose followed by size exclusion chromatography on Superdex to ~95% purity in the Recombinant Protein Core at Northwestern University.

### Binding affinity studies

Binding affinity of different soluble ACE2 variants was examined using an assay which relies on concentration dependence to capture ACE2 enzymatic activity by immobilized SARS-CoV-2 RBD (16). Briefly, purified, his-tagged SARS-CoV-2 S1-RBD protein (1μg/ml) dissolved in Tris-buffered saline (TBS, pH 7.4) was loaded into a 96-well Ni-coated black plate and incubated with shaking for 1 hour at room temperature. After 5 washes with 200μl of TBS supplemented with 0.05% Tween (wash buffer) each, 100μl of soluble ACE2 variants (ACE2 1-618-DDC-ABD, ACE2 1-618-ABD, ACE2 1-618-Fc, ACE2 1-740) dissolved in TBS in concentrations ranging from 0.01ng/ml to 100μg/ml were added and incubated with shaking for 1 hour at room temperature. Wells were then washed 5 times with 200μl wash buffer again, before 90μl of TBS and 10μl of Mca-APK-Dnp substrate were added and fluorescence formation was measured in a microplate fluorescence reader (FLX800, at 320nm using excitation and 400nm emission filters).

### Plasma ACE2 activity after injection of ACE2 1-618-ABD-DDC

Pharmacokinetics of ACE2 1-618-DDC-ABD were examined in male wild-type mice (C57/B6), as previously described (16). For comparison, we also studied native ACE2 1-740. Mice received a single intraperitoneal (i.p.) injection of 1μg/g BW ACE2 1-740 or ACE2 1-618-DDC-ABD. Blood samples were collected in heparinized capillaries by tail-bleeding at different time-points and plasma was isolated by centrifugation. ACE2 enzyme activity was measured using Mca-APK-Dnp substrate (Bachem, Bubendorf, Switzerland).

### Lung ACE2 activity after intranasal administration of ACE2 1-618-DDC-ABD

To demonstrate that intranasal administration of ACE2 1-618-DDC-ABD can increase lung ACE2 activity using a dose comparable to that planned in the *in vivo* studies in k18hACE2 mice, we performed experiments in male wild-type mice (CD1). For lung studies, ACE2 1-618-DDC-ABD was administered intranasally at two different doses (1 and 6-7μg/g BW, both in 40μl PBS). 24 hours later, lungs were harvested. One portion of the lung tissue was snap-frozen at −80°C and the other one was fixed in 10% formalin for subsequent staining studies. The frozen lung tissues were homogenized in RIPA buffer, centrifuged at 8300 RPM for 10 minutes at 4°C and the supernatant saved. In the resulting lung lysates, total ACE2 enzyme activity was measured using Mca-APK-Dnp substrate (Bachem, Bubendorf, Switzerland). For staining studies, an ACE2 antibody AF933 (R&D Systems) was used (16) and staining performed as previously described (32).

### *In vivo* infectivity studies

All work with live SARS-CoV-2 was performed in the BSL-3 facility of the Ricketts Regional Biocontainment Laboratory, operated by The University of Chicago following a protocol approved by the Institutional Animal Care and Use Committees of Northwestern University and University of Chicago. Murine full-length ACE2 binds SARS-CoV-2 poorly (33) and intranasal inoculation of SARS-CoV-2 results in minimal or no clinical disease in wild type mice (28, 34). We therefore used a transgenic mouse that expresses human ACE2, k18-hACE2, and is susceptible for SARS-CoV-2 infection (26–30). K18-hACE2 mice (8 weeks old) were purchased from Jackson. Animals were infected with 2×10^4^ PFU SARS-CoV-2 in 20μl by intranasal inoculation. Mortality in animals infected with this viral load is essentially 100% and animals invariably succumb to disease by days 5-9 (26–29). ACE2 1-618-DDC-ABD (30μl, ~13μg/g BW) was administered intranasally 1 hour prior to viral challenge followed by the same dose intranasally 24 and 48 hours later for a total of 3 doses. Additionally, at the same time-points (1 hour pre- and 24 and 48-hours post-inoculation) an i.p. injection of ACE2 1-618-DDC-ABD (200μl, ~1μg/g BW) was administered. Control animals were treated with the same volumes of PBS intranasally and i.p. performed at the same time-points.

All animals were weighed once a day and monitored twice daily for health using a clinical scoring system (**Table T1**). Animals that lost more than 20% of their baseline body weight or had a clinical score of 3 were sacrificed for humane reasons and this was considered a fatal event. Otherwise, animals that did not meet these criteria were continued to be monitored for up to 14 days while in the BSL-3 facility. Lungs removed from all sacrificed animals were used for viral load measurement by plaque assays (one half), whereas the other lung was fixed in paraffin embedded blocks for histopathology.

### Plaque assay for infectious virus

Tissue samples were collected in DMEM with 2% FBS and were homogenized using 1.4mm ceramic beads in a tissue homogenizer using two 30s pulses. Samples were then centrifuged at 1000g for 5 minutes and the supernatant was serially diluted 10-fold and used to infect VeroE6 cells for 1 hour. Inoculum was removed and 1.25% methylcellulose DMEM solution was added to the cells and incubated for 3 days. Plates were fixed in 1:10 formalin for 1 hour and stained with crystal violet for 1 hour and counted to determine plaque forming units (PFU)/ml.

### Histopathology of the lungs

Formalin-fixed lung sections were embedded in paraffin blocks released from the BSL-3 facility after verifying the absence of infectious virus and used to generate slides for staining studies by the Mouse Histology & Phenotyping Laboratory (MHPL) center, Northwestern University, Chicago. Histopathology was evaluated by two expert lung pathologists. One blinded lung pathologist evaluated the severity and presence of lung injury using a scoring system recently described in k18-hACE2 mice infected with SARS-CoV-2 (26). The alterations scored were mononuclear infiltrates, alveolar hemorrhage, edema, cellular necrosis, hyaline membranes and thrombosis. The scale was as follows: 0 = no detection, 1 = uncommon detection in <5% lung fields (200Å~), 2 = detectable in up to 30% of lung fields, 3 = detectable in 33-66% of lung fields and 4 = detectable in >66% of lung fields. Neutrophil infiltration was evaluated on a scale of 0-3 as follows: 0 = within normal range, 1 = scattered PMNs sequestered in septa, 2 = score 1 and solitary PMNs extravasated in airspaces, 3 = score 2 plus and aggregates in vessel and airspaces.

## Results

### Generation of a soluble ACE2 variant with enhanced binding to the receptor binding domain of SARS-CoV-2

A cDNA coding for the C-terminal portion of a 618 amino acids fragment of human ACE2 protein (termed *ace2 1–618*) was fused with the *abd* cDNA encoding for a small ABD protein (5-kD) as recently described by us (16). The dimerization of the ACE2 1-618-ABD chimeric protein was achieved by inserting a cDNA coding for a hinge-like region containing a dodecapeptide (DDC) between *g4s3* and the c-terminus of the cDNA for *ace2 1-618* (**figure S1**). After verifying the DNA sequence of the pcDNA3-4 fused with the cDNA for *ace2 1–618-ddc-abd*, the protein was produced in expi293 cells and purified to ~95% purity.

The binding of ACE2 1-618-DDC-ABD to the SARS-CoV-2 S1-RBD was compared to other soluble ACE2 proteins including ACE2 1-618-ABD and the native ACE2 1-740 protein. The binding affinity of ACE2 1-618-DDC-ABD was higher than that of native ACE2 1-740 and ACE2 1-618-ABD **(figure 1A)**. This is also evident from the half maximal effective concentration (EC50) which was significantly lower for ACE2 1-618-DDC-ABD than for ACE2 1-740 (p=0.0223) and much lower than for ACE2 1-618-ABD (p=0.0039), calculated by repeated measures analysis.

**Figure 1.**
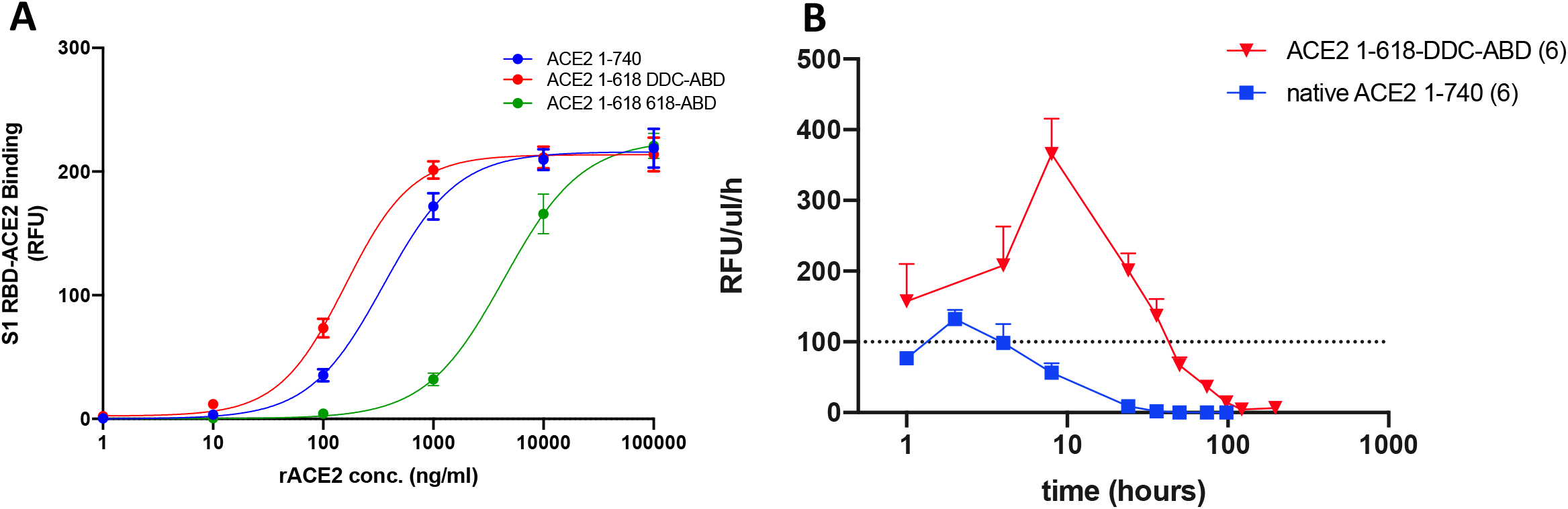
Binding affinity to the Receptor Binding Domain (RBD) of the viral S1 Glycoprotein (Panel A) and pharmacokinetics of ACE2 1-618-DDC-ABD protein (Panel B) as compared to native soluble ACE2. **Panel A.** The binding of ACE2 variants 1-618-DDC-ABD (red), 618-ABD (green), and the native ACE2 740 protein (blue) to SARS-CoV-2 S1-RBD was studied using a recently published assay (16). The binding affinity of the ACE2 variant 1-618-DDC-ABD is markedly higher as compared to that of the ACE2 1-618-ABD and also slightly higher than that of native ACE2 740 soluble protein. **Panel B.** The ACE2 1-618-DDC-ABD (red) variant resulted in higher plasma ACE2 activity than native soluble ACE2 1-740 (blue) post intraperitoneal injection. Moreover, after native ACE2 1-740 administration, plasma ACE2 activity decreased already after 8 hours post injection whereas ACE2 618-DDC-ABD resulted in markedly increased duration of action as shown by substantial plasma ACE2 activity at 72 hours.

### Prolonged duration of plasma ACE2 enzymatic activity after injection of ACE2 1-618-DDC-ABD

We then examined the *in-vivo* plasma enzymatic activity of the ACE2 1-618-DDC-ABD variant as a measure of protein duration of action and compared it to the native ACE2 1-740 injected also intraperitoneally (**figure 1B**). By measuring plasma enzymatic activity, one cannot discriminate between exogenous human plasma activity (from the infused protein) and endogenous mouse ACE2 activity. Baseline plasma ACE2 activity in mice, however, is usually very low (23, 35). Therefore, the observed activity after injection of soluble ACE2 proteins essentially reflects exogenous activity from the injected soluble ACE2 protein. After injection of ACE2 1-618-DDC-ABD, plasma ACE2 activity peaked at 8 hours (365 RFU/μl/hr) with substantial plasma ACE2 activity remaining at 24 hours (200 RFU/μl/hr), 36 hours (136 RFU/μl/hr), and 48 hours (66 RFU/μl/hr) (**figure 1B**). Plasma ACE2 activity was still substantial at 72 hours (36 RFU/μl/hr) post injection. By contrast, after injection of the native ACE2 1-740, plasma activity had decreased markedly by 8 hours (56 RFU/μl/hr) and was reduced to very low levels thereafter (**figure 1B**).

### Lung ACE2 activity after intranasal delivery

To demonstrate that intranasal injection of ACE2 1-618-DDC-ABD results in lung uptake presumably in the alveolar space we measured ACE2 protein presence in the lungs. ACE2 protein levels in lungs of mice is essentially undetectable by Western Blot and enzymatic activity likewise is very low (36–39). 24 hours after intranasal administration of ACE2 1-618-DDC-ABD (1 or 6-8μg/g BW) we were able to detect variable but substantial ACE2 activity in the lung of 5 out of 6 injected mice but not in non-infused animals (**figure S2A**). Staining of the lung with an ACE2 antibody, confirmed the presence of ACE2 in the lung of animals infused with ACE2 1-618-DDC-ABD as compared to non-infused animals that had barely detectable staining for ACE2 (**figure S2B**).

### ACE2 1-618-DDC-ABD administration to k18-hACE2 male and female mice infected with SARS-CoV-2

Since mice are resistant to SARS-CoV-2 infection we used the transgenic k18-hACE2 mouse that is susceptible to SARS-CoV-2 infection (25–30). After SARS-CoV-2 inoculation with 2×10^4^-2×10^5^ PFU these animals have essentially 100% mortality by days 5-9 (26–29). We used the same lethal dose as in these previous studies. In total, 20 animals (10 male, 10 female) were infected with SARS-CoV-2. Either ACE2 1-618-DDC-ABD or PBS (n=10 per group) was administered intranasally under light anesthesia 1 hour before virus inoculation and 24 and 48 hours later. In addition to the 3 intranasal doses, ACE2 1-618-DDC-ABD or PBS vehicle were administered i.p. at the same three time points.

The 5 control female mice, PBS treated, lost more than 20% of body weight on day 6. By contrast, the ACE2 1-618-DDC-ABD treated mice had only minimal weight loss by day 6 (**figure 2A**). By day 6, all the 5 female PBS-treated also had a worsening clinical score whereas in the protein treated female animals the clinical score never went up above 2 and returned to a normal score of 1 after day 6 (**figure 2B**). By study protocol, the animals that lost more than 20% body weight were considered to have reached the mortality end-point and were euthanized for humane reasons. Therefore, mortality rate by day 6 was 100% in PBS-treated and 0% in ACE2 1-618-DDC-ABD treated female mice (**figure 2C**).

**Figure 2.**
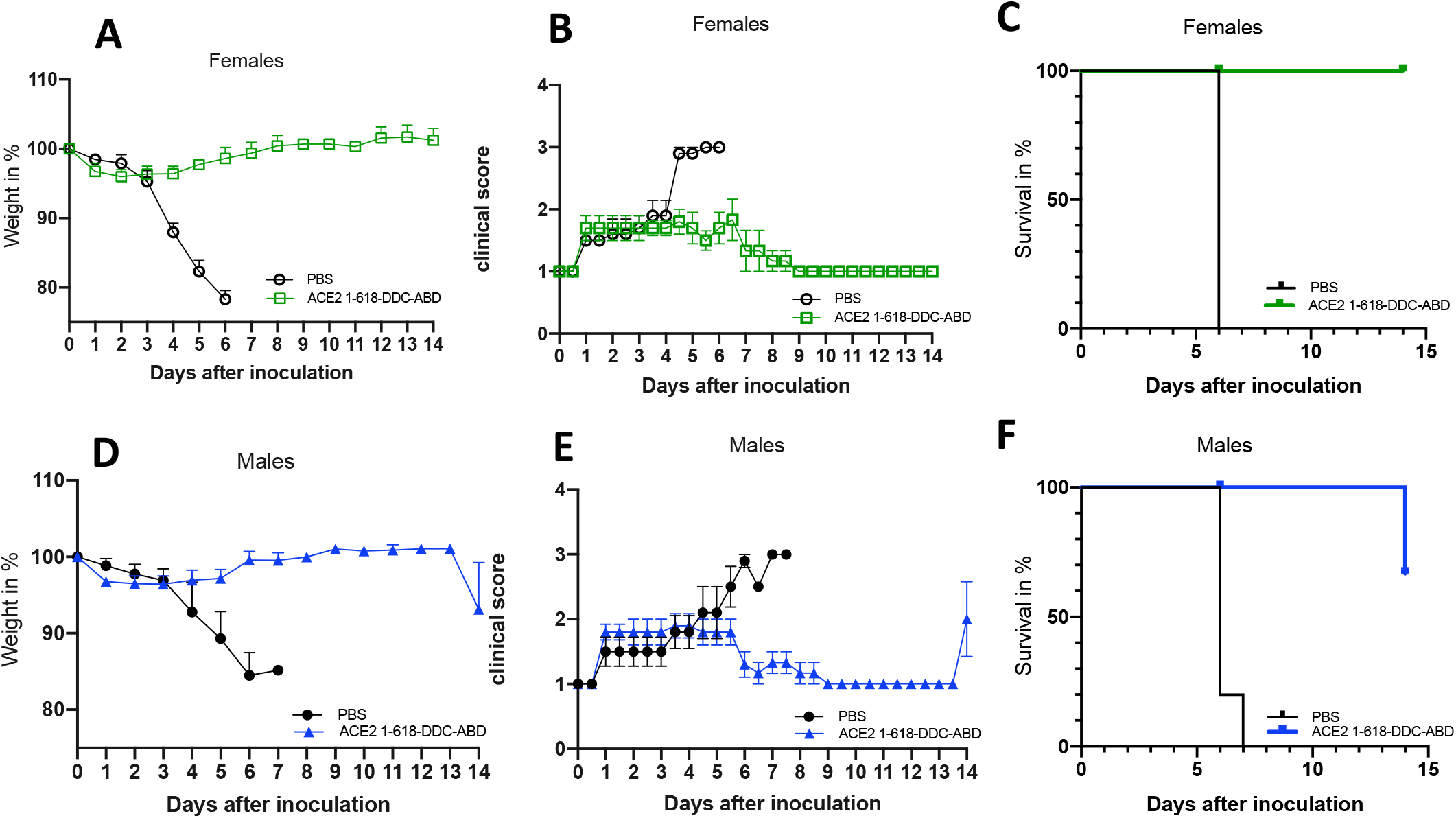
Prevention of mortality and improvement of clinical parameters after ACE2 1-618-DDC-ABD administration to female and male k18-hACE2 mice infected with SARS-CoV-2. The administration of ACE2 1-618-DDC-ABD largely prevented the body weight loss (**Panels A and D**), improved clinical scores (**Panel B and E**) and prevented mortality (**Panels C and F**) in five k18-hACE2 transgenic female (upper panels) and five male mice (lower panels) as compared to vehicle-mice (PBS) infected with SARS-CoV-2 (also 5 male and 5 female mice).

Of the 5 control male mice, vehicle-treated, 2 animals lost more than 20% of body weight by day 6; the remaining 3 also had lost weight but less than 20% body weight (11, 15 and 16%) (**figure 2D**). They also had to be sacrificed by study protocol on day 6 (2 of them) and day 7 (the remaining one) because they were very sick by clinical score (**figure 2E**). Therefore, all 5 male untreated mice reached the mortality endpoint by day 6/7 (**figure 2F**). By contrast, the 5 male ACE2 1-618-DDC-ABD treated mice experienced a small transient weight loss (**figure 2D**) and their mean clinical score was less than 2 up to day 6 (**figure 2E**). After day 6, the treated male mice all returned to a normal score of 1 (**figure 2E**). Accordingly, at this time point there was no need to sacrifice any of the male treated animals with ACE2 1-618-DDC-ABD and therefore the mortality was 0% (**figure 2F**).

To be able to have viral titers at the same time as the untreated animals we sacrificed 2 males and 2 females in the treated group on day 6. The remaining male and female mice were monitored for an additional week during which time they had a completely normal clinical score (**figure 2B and E**) and stable weight (**figure 2A and D**). One of the protein-treated male mice that was deemed to be healthy by clinical score, but by day 14, lost 20% of its body weight and became sick (**figure 2D and E**). This male mouse was euthanized together with all the remaining male and female mice that were healthy at this time point. This male was considered a mortality event on day 14 (**figure 2F**).

### Viral titers are decreased in lungs of mice treated with ACE2 1-618-DDC-ABD

Administration of ACE2 1-618-DDC-ABD resulted in a marked reduction of SARS-CoV-2 lung titers measured by plaque assay in k18-hACE2 mice. At 6 days post infection (except for a male mouse that was not euthanized until day 7 per study protocol), titers were high ranging from 10^2^ to 10^7^ PFU/ml in all control PBS treated animals. By contrast, at the same time point (day 6) in 2 males and 2 females treated with ACE2 1-618-DDC-ABD, lung viral titers were in the low range in 2 animals and undetectable in the other 2 (**figure 3**). In the 6 remaining k18-hACE2 mice that had received ACE2 618-DDC-ABD and were sacrificed 14 days after viral inoculation, no viral titers were detectable in any of them when they were sacrificed (**figure 3**).

**Figure 3.**
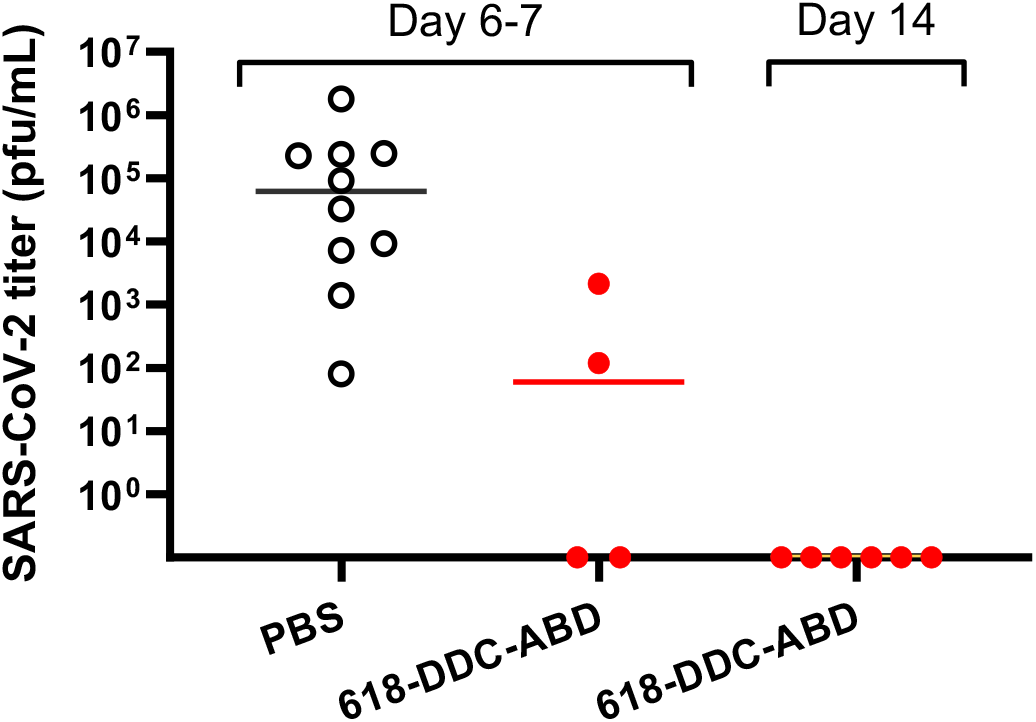
Administration of ACE2 618-DDC-ABD to k18-hACE2 mice infected with SARS-CoV-2 resulted in reduction of SARS-CoV-2 lung titers measured by plaque assay. At 6-7 days post infection, titers were high in all untreated animals (PBS, black) that had to be humanely euthanized by study design. In contrast, at the same time point (day 6) in 2 male and 2 female ACE2 618-DDC-ABD treated mice (red) that were healthy but were sacrificed, lung viral titers were lower or undetectable. No SARS-CoV-2 virus was detectable in the remaining k18-hACE2 mice that received ACE2 618-DDC-ABD and were sacrificed 14 days after viral inoculation.

### Histopathology is improved in animals treated with ACE2 1-618-DDC-ABD

Examination of a lung from each one of the 10 untreated animals showed extensive cellular infiltrates evident at low magnification (**figure S3, left panels**). The lesions consistently seen in untreated animals were dense perivascular mononuclear infiltrates and scattered neutrophils (**figure 4A**) with alveolar edema and hemorrhage (**figure 4B**). In sharp contrast, the lungs of treated animals sacrificed at the same time as the untreated ones (day 6) showed much less cell infiltrates and only mild to moderate alveolar hemorrhage (**figure 4A and B, S3 and S4 right panels**). In treated animals sacrificed at day 14, the lesions were further improved and in some cases the lungs appeared normal (**figure 4A and B, S3 and S4, right panels**).

**Figure 4.**
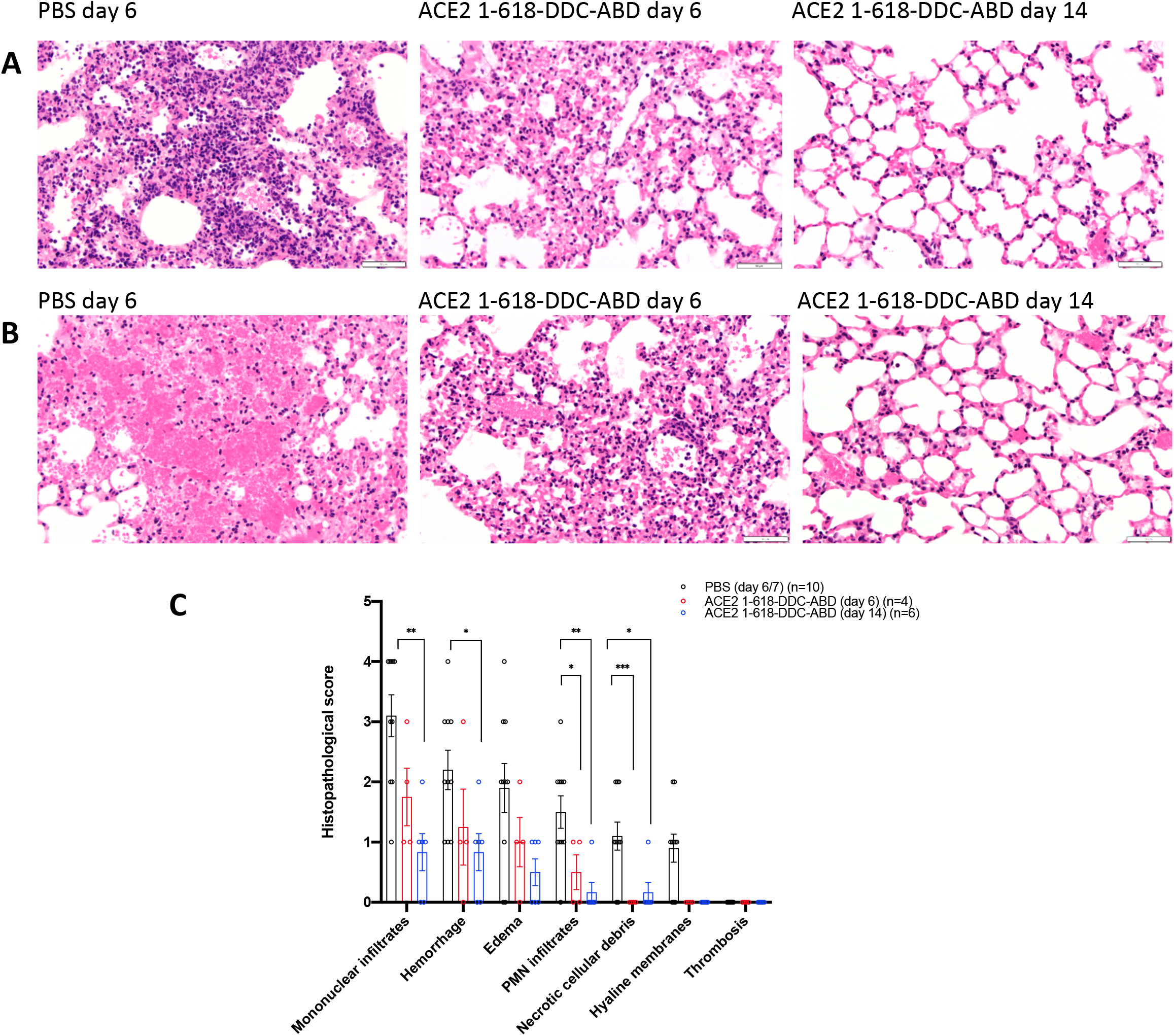
Representative example of lung histopathology (Panel A and B) and lung injury scores (Panel C). **Panel A.** Representative lung histopathology in k18-hACE2 mice infected with SARS-CoV-2 on day 6 (left and middle) and day 14 (right). Untreated animals (PBS-vehicle) show dense perivascular mononuclear infiltrates and scattered neutrophils (left). **Panel B.** Untreated animals (PBS-vehicle) show more extensive alveolar hemorrhage (left) in contrast to animals treated with ACE2 1-618-DDC-ABD on days 6 and 14. Hematoxylin-Eosin, original magnification X400. **Panel C.** The histopathological lung injury scores are lower in animals treated with ACE2 1-618-DDC-ABD both on day 6 (red) and day 14 (blue) than vehicle treated controls on day 6/7 (black). Significance is indicated in the figure (*** = p < 0.001; ** = p < 0.01; * = p < 0.05) was calculated using ANOVA followed by post-hoc Dunnett’s multiple comparisons test.

To semi-quantitate the findings, we used a scoring system recently used in the same mouse transgenic model infected with SARS-CoV-2 (26) (**figure 4B**). All untreated animals had mononuclear infiltrates (mean 3.1 ±0.3), alveolar hemorrhage (mean 2.2 ±0.3), edema (mean 1.9 ±0.4) and neutrophils (mean 1.5 ±0.3). In contrast, treated animals sacrificed at day 6 had lower scores for mononuclear infiltrates (mean 1.75 ±0.3), alveolar hemorrhage (mean 1.25 ±0.4), edema (mean 1 ±0.3) and neutrophils (mean 0.5 ±0.2). The scores of lungs from animals sacrificed at day 14 were lower than in the other 2 groups; mononuclear infiltrates (mean 0.8 ±0.2), alveolar hemorrhage (mean 0.8 ±0.2), edema (mean 0.5 ±0.2) and neutrophils (mean 0.16 ±0.1). The differences were all significant as indicated in the legend of figure 4. Necrotic cellular debris was seen in lungs of a few untreated animals and only one treated animal at 14 days. Hyaline membranes were seen also in a few untreated animals but none of the treated animals. Clear-cut thrombosis was not seen in untreated or treated animals (**figure 4B**).

## Discussion

In this study we used a novel dimeric construct of a truncated and longer acting ACE2, termed ACE2 1-618-DDC-ABD, that displayed an almost 30-fold enhanced affinity for SARS-CoV-2 S1-RBD, compared with its monomeric version (ACE2 1-618-ABD) lacking only the disulfide crosslinking hinge peptide (**figure 1**). Our findings demonstrate for the first-time complete efficacy *in vivo* of a soluble ACE2 protein for the prevention/ treatment of SARS-CoV-2 induced disease in a mouse model susceptible to infectivity by this coronavirus. Since mice are resistant to SARS-CoV-2 infection, we use k18-hACE2 mice expressing human ACE2 that have essentially 100% mortality when infected with a high viral dose of SARS-CoV-2 (26–30). In this model, the efficacy of a novel soluble human ACE2 protein was demonstrated by reducing mortality from 100% to 0% in both male and female animals at day 6 post viral inoculation (**figure 2**). ACE2 1-618-DDC-ABD administration resulted in prevention of severe weight loss and marked improvement in clinical scores (**figure 2**). These improvements were associated with reduced lung viral titers of SARS-CoV-2 by plaque assay (**figure 3**) and marked improvements of lung histopathology (**figure 4**). Specifically, examination of lungs from untreated, infected mice showed extensive damage mainly consisting of alveolar hemorrhage, perivascular infiltration and interstitial edema. The infiltrates consisted mainly of mononuclear cells, but neutrophils were also seen frequently. In sharp contrast, the lungs of treated animals showed mild to moderate damage 6 days post infection and a marked improvement 14 days post infection (**figure 4A**).

The design of this study was in part preventative as the soluble ACE2 protein was given 1 hour prior to viral inoculation and then continued daily for 2 days. We chose this approach as proof of concept that our soluble human ACE2 protein can obliterate SARS-CoV-2 infectivity at least when given early following viral exposure. The rationale for the potential effect of soluble ACE2 protein for treatment and prevention of SARS-CoV-2 infection was outlined by us (12) and others in early 2020 (40). That is, provision of sufficient amounts of soluble ACE2 protein likely acts as a decoy to bind to the spike of SARS-CoV-2 RBD and therefore should limit viral uptake mediated by membrane-bound full-length ACE2 (**figure 5**). Consequently, SARS-CoV-2 entry into the cells and viral replication should be prevented. In addition, with more advanced disease, further therapeutic benefit should result from restoring decreased ACE2 activity and therefore, correction of the altered balance of Angiotensin II and bradykinins resulting from deficiency of ACE2 enzymatic activity (11). ACE2 degrades not only Angiotensin II but also the pro-inflammatory peptide des-arg^9^ bradykinin (41–43). ACE2, therefore, acts as a guardian by preventing excessive levels of peptides with pro-inflammatory effects and protects the lungs from local injury (11, 42, 44). ACE2 enzymatic activity, therefore, is an important feature that should be retained in ACE2 proteins to prevent and treat SARS-CoV-2 infection. Membrane bound full-length ACE2 is believed to be deficient because of internalization of the ACE2-SARS-CoV-2-complex which, in turn, contributes to lung injury owing to accumulation of Angiotensin II and des-Arg^9^ Bradykinin deficiency (11, 40, 44–47).

**Figure 5.**
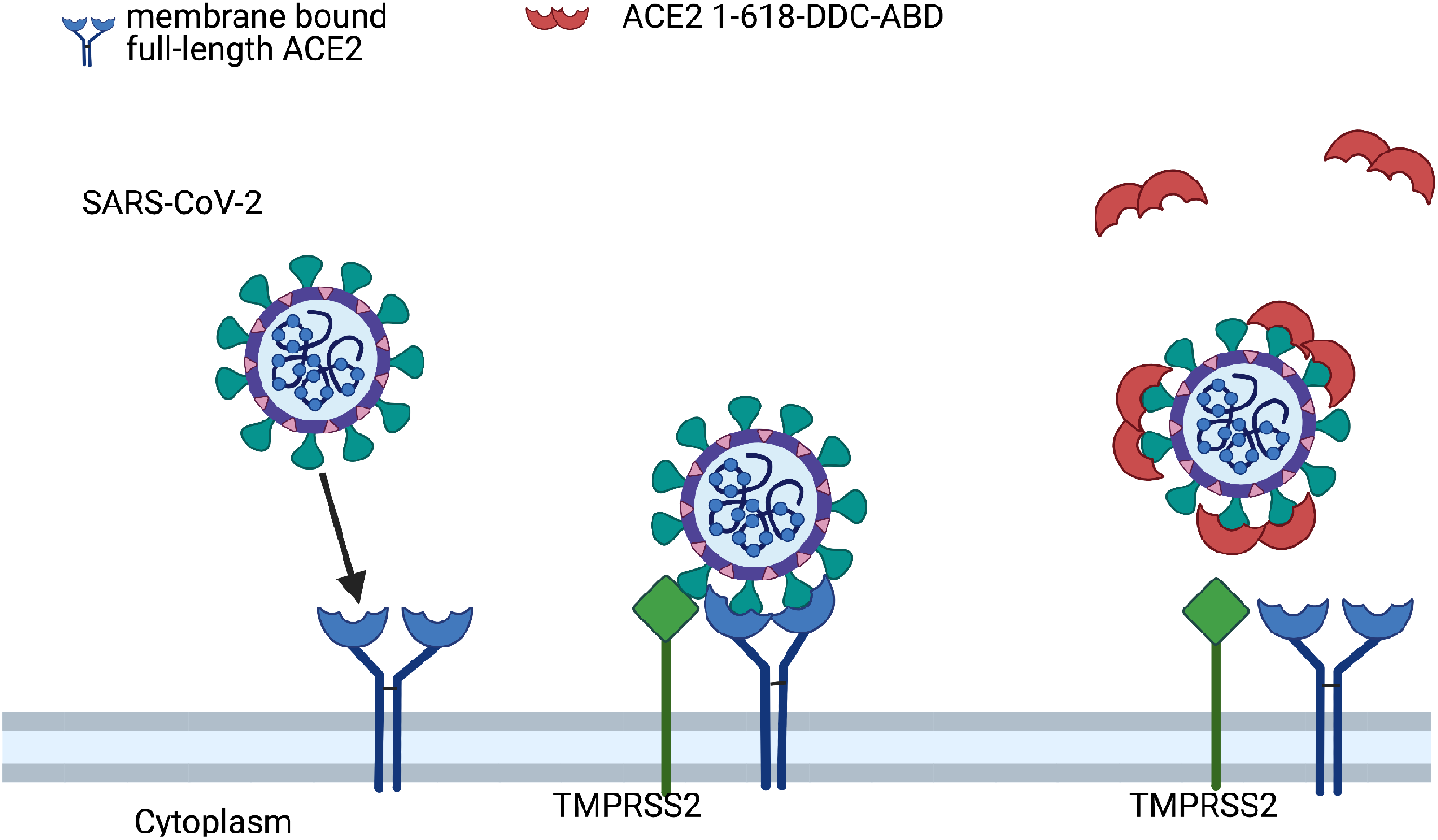
Postulated mechanism of action of ACE2 1-618-DDC-ABD. Administered ACE2 1-618-DDC-ABD (red semi-circles) binds to SARS-CoV-2 acting as a decoy to prevent the binding of SARS-CoV-2 to membrane bound full-length ACE2 receptors (blue). This prevents the internalization of the ACE2-SARS-CoV-2 complex activated by TMPRSS2 (green). Modified from Davidson et al, Hypertension 2020 (10). Created with biorender.com.

There have been recent reports on soluble ACE2 proteins that tested their decoy effect *in vitro* (13–20). To date, to our knowledge, there is only one report testing soluble ACE2 variants for SARS-CoV-2 infection in an animal model. This pre-print from July 2020, which is still unpublished, reported an ACE2-Fc variant with a mutated Fc-region to inhibit binding to the Fc-receptor and also lacked ACE2 enzymatic activity (15). This variant was tested in a mouse model made susceptible to SARS-CoV-2 infection by inoculation with adenovirus encoding human ACE2 (15). Staining with SARS-CoV-1 antiserum by immunofluorescence of the lung was reported to be decreased in the group treated with the ACE2-Fc variant (15). Mortality, lung histology and other clinical parameters, however, were not reported (15). ACE2 proteins tagged with Fc also have the advantage of extended duration of action (15, 17, 20, 48). It has been noted, however, that Fc-receptor activation could potentially lead to complement dependent cytotoxicity, antibody dependent cytotoxicity and infection via the CD16 Fc receptor (49). To avoid these uncertain effects, our approach was to further modify a shorter ACE2 variant fused with an albumin binding domain (ABD) tag to extend its duration of action (16). This fused protein was shown to have a duration of action of at least 3 days and to also neutralize SARS-CoV-2 in human kidney organoids (16). To improve the binding capacity to the SARS-CoV-2 RBD, we fused this protein with a hinge like region termed DDC (31). This resulted in dimerization (**figure S1**) which confers a marked increase in binding to the RBD of SARS-CoV-2 as compared to ACE2 1-618-ABD; it also higher binding than the native soluble ACE2 1-740 (**figure 1A**). We think this feature renders the ACE2 1-618-DDC-ABD variant much improved in that the dosing required to intercept SARS-CoV-2 could be reduced owing to the enhanced binding characteristics while retaining the prolonged duration of action conferred by the ABD fusion.

There are advantages of administering a soluble ACE2 protein that acts as a decoy to intercept SARS-CoV-2 from binding to the cell membrane bound ACE2 receptor. Soluble ACE2 can be administered preventatively to those at risk for severe COVID-19 infection or recently infected individuals, whereas convalescent plasma and antibody infusions are not designed for preventative use. Mutations of SARS-CoV-2, that are increasingly recognized (50–52), moreover, might escape antibody-based therapies. Soluble ACE2, by contrast, ought to be effective in neutralizing mutated variants of SARS-CoV-2 with the rare exception of a mutation that would happen to interfere with the RBD of SARS-CoV-2 for the soluble ACE2 protein that is administered. Such unlikely mutation, of course, would be a blessing to humanity since it would also prevent the binding to the cell membrane bound FL-ACE2 and therefore cell entry and replication. The preventative effect of our approach, moreover, is exerted rapidly, while the vaccination derived immunity via antibodies needs longer to reach a protective level. Therefore, in people exposed to SARS-CoV-2 who for whatever reason were not vaccinated, the use of ACE2 1-618-DDC-ABD as an inhalant (see below) could provide immediate protection.

It is important to consider the route of administration. In our study, we gave ACE2 1-618-DDC-ABD both intranasally and systemically and therefore cannot distinguish with certainty which venue was more effective in neutralizing SARS-CoV-2. Considering that the nasal and lung epithelia are considered the entry sites for SARS-CoV-2 it is likely that the intranasal route exerted the bulk of the protective effect. The presence of ACE2 activity, normally is barely detectable in the lung, where ACE2 is restricted to type 2 pneumocytes (36–39). After intranasal administration to uninfected control mice our pilot studies revealed detectable ACE2 activity and protein staining (**figure S2**). We think that intranasal is the preferred route of administration. In our design, however, we wanted to cover the possibility of systemic effects and brain invasion that occurs in some k18-hACE2 mice infected with SARS-CoV-2 (26–29). Accordingly, we also gave a concurrent systemic dose i.p. for 3 days. Studies using the soluble ACE2 protein only intranasally or systemically will need to be done to address the issue of route of administration. Dosing studies will also be needed since we chose a high dose in this initial preclinical study to demonstrate efficacy beyond any doubt. Finally, it should be noted that the enzymatic activity of our soluble protein will likely add to a therapeutic action by restoring ACE2 activity in the lungs which is likely depleted when SARS-CoV-2 ACE2 complexes are formed and internalized (11, 44). In the process the ability to metabolize substrates of ACE2 such as Angiotensin II and des-Arg bradykinin are compromised and these peptides can promote lung injury (42, 44, 46). This enzymatic action provides, therefore, a therapeutic component of soluble ACE2 proteins that retain enzymatic activity.

It is worthy to note, that, to our knowledge, this is the first time that a protein that presumably uses the decoy mechanism to neutralize a virus is fully effective *in vivo*. There are several examples *in vitro*, of compounds that use the decoy mechanism to inhibit cell entry of dengue virus. Heparin has been used in human liver cell lines to compete with heparin sulfate, the cell entry receptor for dengue virus (53). The glycosaminoglycan-mimetic Suramin showed an antiviral effect on the human adenovirus D37 (HAdV-D37) that causes epidemic keratoconjunctivitis (54). Glycosaminoglycans are the cell entry receptor for HAdV-D37, and Suramin inhibited the attachment of HAdV-D3 to human corneal epithelial cells (54). Our findings that ACE2 1-618-DDC-ABD can neutralize SARS-CoV-2 *in vivo* support further research with other compounds that may use the decoy effect therapeutically more broadly.

In summary, the administration of a novel soluble human ACE2 variant with extended duration of action and increased binding efficacy for the SARS-CoV-2 RBD to transgenic k18-hACE2 mice, infected with SARS-CoV-2, resulted in complete resolution of symptoms, marked improvement of lung histopathology and prevented the universal lethality observed in untreated animals. Strategies using the same approach are quite likely to be effective in the prevention and treatment of COVID-19 and become an essential complement to combat the present pandemic and future outbreaks of other coronavirus that use ACE2 as its main receptor.

## Author contributions

L. Hassler performed several experiments and helped with the writing of the manuscript and data analysis. J. Wysocki designed and performed many of the experiments and was involved with the writing of the paper. I. Gelarden and A. Yeldandi evaluated the lung histology. G. Randall supervised the infectivity studies in the Biosafety Level 3 facility performed by A. Tomatsidou and V. Nicoleascu. J. Henkin was involved in the writing of the manuscript and analysis of the findings. D. Batlle supervised the overall project, designed experiments and wrote most of the paper.

## Acknowledgments

We are very grateful for Dr. Dominique Missiakas, Howard T. Ricketts Laboratory Director, for coordinating the studies performed in the BSL-3 facility of the Ricketts laboratory and Dr. Sergii Pshenychnyi, managing director at the recombinant Protein Production Core (rPPC), for supervising the production and purification of the ACE2 1-618-DDC-ABD variant in the recombinant Protein Production Core at Northwestern University, Evanston. We also acknowledge the George M. O’Brien Kidney Research Core Center (NU GoKidney) supported by the award P30 DK114857 (National Institute of Diabetes and Digestive and Kidney Diseases). We also acknowledge the support of a gift from the Joseph and Bessie Feinberg Foundation and a gift from the state of Dr. Frank Krumlowsky.

