## Supplement for "A novel soluble ACE2 protein totally protects from lethal disease caused by SARS-CoV-2 infection"

### Short title: Novel soluble ACE2 to combat SARS-CoV-2

Luise Hassler[1], Jan Wysocki[1], Ian Gelarden[1], Anastasia Tomatsidou[2,3], Haley Gula[2,3], Vlad Nicoleascu[2,3], Glenn Randall[2,3], Jack Henkin[4], Anjana Yeldandi[1], and Daniel Batlle[1]

[1] From Division of Nephrology/Hypertension, Department of Medicine and the Department of Pathology, Northwestern University, Feinberg School of Medicine, Chicago, Illinois

[2] Department of Microbiology, The University of Chicago, Chicago, Illinois

[3] Ricketts Regional Biocontainment Laboratory, University of Chicago, Lemont, Illinois

[4] Center for Developmental Therapeutics, Northwestern University, Evanston, IL, USA

**Correspondence:**

Dr. Daniel Batlle

Division of Nephrology and Hypertension, Department of Medicine

Northwestern University, The Feinberg School of Medicine

320 E Superior, Tarry 14-727, Chicago, IL 60611

**Table of content:**

- Table T1
- Supplementary figures S1-S4

**Table T1:** Scoring system in BSL-3 facility for health evaluation of mice infected with SARS-CoV-2.

| Score | Description |
|---|---|
| 0 | Pre-inoculation: mice are bright, alert, active, normal fur coat and posture |
| 1 | Post-inoculation (PI): mice are bright, alert, active, normal fur coat and posture, no weight loss |
| 1.5 | Mice present with slightly ruffled fur but are active, or weight loss might occur but <2.5%, recovery can be expected |
| 2 | Ruffled fur or less active or weight loss <5%, recovery might occur |
| 2.5 | Ruffled fur or not active but moves when touched or hunched posture or difficulty breathing or weight loss 5-10%, Recovery is unlikely but still might occur |
| 3 | Ruffled fur or inactive but moves when touched or difficulty breathing or weight loss at 11-20%, recovery is not expected |
| 4 | Ruffled fur or positioned on its side or back or dehydrated or difficulty breathing or weight loss >20% or labored breathing, recovery is not expected |
| 5 | death |

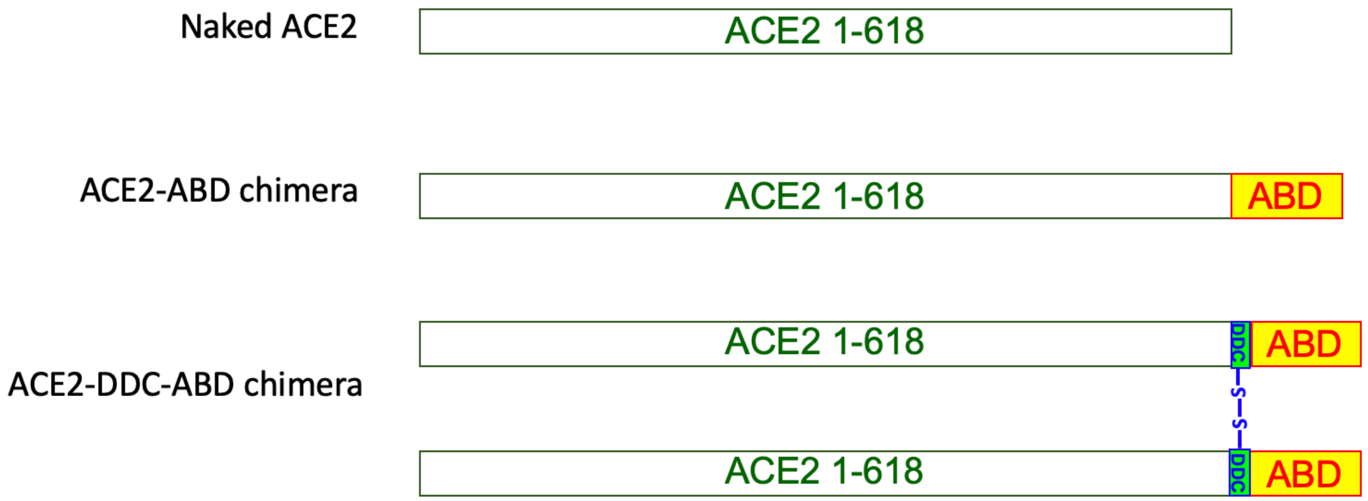


**Figure S1. Scheme of soluble ACE2 1-618 variants.**

The upper panel shows naked ACE2 1-618, a monomer with duration of action of only few hours. The middle panel shows ACE2 1-618-ABD, which was tagged with an Albumin Binding Domain (ABD) to extend its duration of action to days. The lower panel shows the scheme of ACE2 1-618-DDC-ABD, which is a chimera of ACE2 1-618-ABD fused with a hinge-like region containing a dodecapeptide (DDC) for dimerization and increased binding to the receptor binding domain of SARS-CoV-2 (see text).

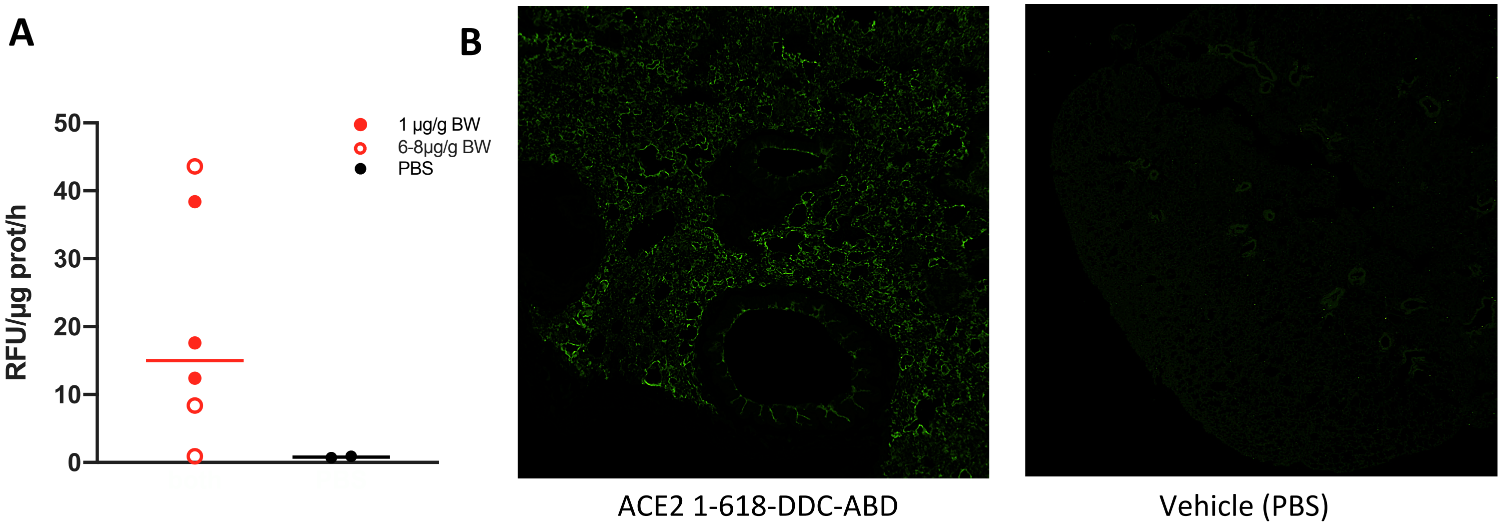


**Figure S2. Lung ACE2 activity and staining 24 hours after intranasal administration of ACE2 1-618-DDC-ABD.** ACE2 1-618-DDC-ABD (1 or 6-8µg/g BW) or PBS was administered intranasally to wild-type mice and lung ACE2 activity (**Panel A**) and staining for ACE2 (**Panel B**) was performed 24 hours later. Animals that received ACE2 1-618-DDC-ABD had variable but substantial lung ACE2 activity whereas controls did not (**Panel A**). Staining of the lung with an ACE2 antibody (**Panel B**) confirmed the presence of ACE2 in alveoli (left) compared to a control that received vehicle (PBS) (right).

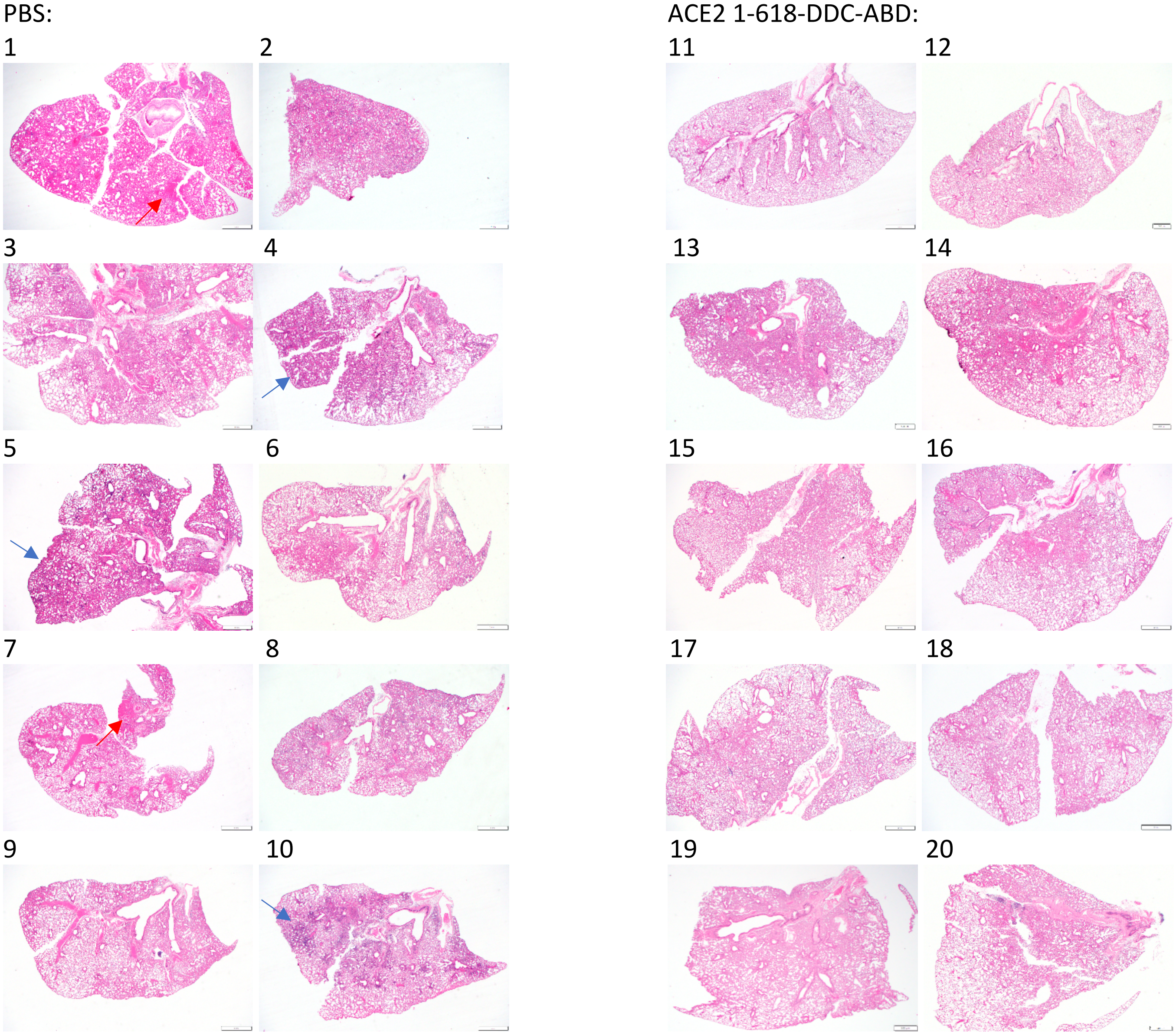


**Figure S3. Histopathology of lungs stained with hematoxylin and eosin, original magnification X20.**

An overview of sections of one lung from SARS-CoV-2 infected animals untreated (n=10, left panel) or treated with ACE2 1-618-DDC-ABD (n=10, right panel). Overall, lungs of all untreated animals show extensive perivascular and interstitial basophilic mononuclear infiltrates (blue arrows show representative foci) and eosinophilic alveolar hemorrhage (red arrows show representative foci) compared to treated animals with much less apparent involvement and in some cases appear essentially normal at low power (cases 11, 12, 15, 17, 18, and 19). Cases 1-10 (left) are untreated and sacrificed at day 6/7 post viral inoculation, cases 11-14 (right) are treated with ACE2 1-618-DDC-ABD and sacrificed on day 6, cases 17-20 (right) are also treated and were sacrificed on day 14 post viral inoculation.

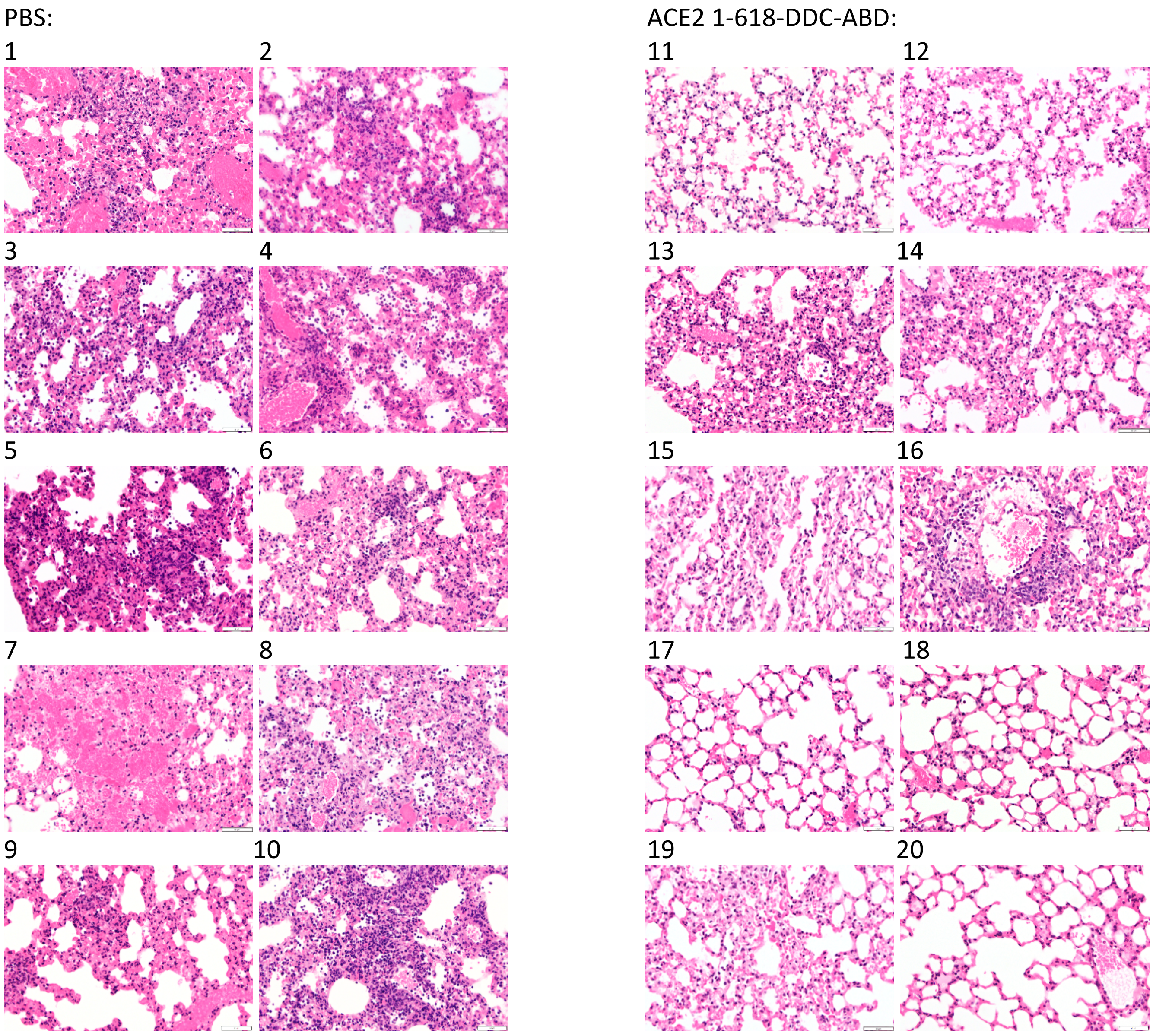


**Figure S4. Histopathology in lungs stained with hematoxylin and eosin,** original magnification X400. Examination of lungs showed extensive alveolar damage with hemorrhage, perivascular and interstitial infiltrates, in all untreated animals (left panels). In sharp contrast, the lungs of animals treated with ACE2 1-618-DDC-ABD (right panels) show less alveolar damage, with many cases appearing largely normal (11, 12, 15, 17, 18, 19, and 20). Cases 1-10 (left) are untreated and sacrificed at day 6/7 post viral inoculation, cases 11-14 (right) are treated with ACE2 1-618-DDC-ABD and sacrificed on day 6, cases 17-20 (right) are also treated and were sacrificed on day 14 post viral inoculation.
